# Virus-like particle-based vaccines targeting the *Anopheles* mosquito salivary protein, TRIO

**DOI:** 10.1101/2024.09.05.611467

**Authors:** Alexandra Francian, Yevel Flores-Garcia, John R. Powell, Nikolai Petrovsky, Fidel Zavala, Bryce Chackerian

## Abstract

Malaria is a highly lethal infectious disease caused by *Plasmodium* parasites. These parasites are transmitted to vertebrate hosts when mosquitoes of the *Anopheles* genus probe for a blood meal. Sporozoites, the infectious stage of *Plasmodium*, transit to the liver within hours of injection into the dermis. Vaccine efforts are hindered by the complexity of the parasite’s lifecycle and the speed at which the infection is established in the liver. In an effort to enhance immunity against *Plasmodium*, we produced a virus-like particle (VLP)-based vaccine displaying an epitope of TRIO, an *Anopheles* salivary protein which has been shown to enhance mobility and dispersal of sporozoites in the dermis. Previous work demonstrated that passive immunization with TRIO offered protection from liver infection and acted synergistically with a *Plasmodium* targeted vaccine. Immunization of mice with TRIO VLPs resulted in high-titer and long-lasting antibody responses that did not significantly drop for over 18 months post-immunization. TRIO VLPs were similarly immunogenic when combined with an anti-malaria vaccine targeting the L9 epitope of the *Plasmodium falciparum* circumsporozoite protein.However, when used in a malaria challenge mouse model, TRIO VLPs only provided modest protection from infection and did not boost the protection provided by L9 VLPs.

## INTRODUCTION

Malaria remains a major global public health concern and one of the most lethal infectious diseases worldwide. According to the latest report from the World Health Organization (WHO), there were an estimated 249 million malaria cases and 608,000 deaths in 2022^1^. Although multiple species of the *Plasmodium* parasite can cause malaria, *Plasmodium (P*.*) falciparum* is responsible for the most severe form of the disease with the highest morbidity and mortality, especially impacting children under 5 years old. Infection begins when an *Anopheles* mosquito injects *P. falciparum* sporozoites while probing for a blood meal. These sporozoites migrate from the skin to the liver, where they multiply and produce merozoites that trigger the symptoms of malaria upon infecting erythrocytes^2^. The need to develop new approaches to control malaria becomes even more urgent with the rise of drug-resistant parasites and insecticide-resistant mosquitoes, as well as the expanding range of vector species due to increased urbanization and climate change^3,4^. Two recently approved vaccines, RTS,S/AS01 and R21/Matrix-M, offer protection from infection, but the challenge of waning vaccine-induced immunity underscores the need for additional strategies to augment immunity against *P. falciparum*^5,6^.

Next generation vaccines targeting different stages of the *P. falciparum* lifecycle, including the pre-erythrocytic stages, blood stage, and transmission stage, are currently in development.While each vaccine class has potential, the most successful vaccines to date target the pre-erythrocytic parasite, due to the possibility of preventing initial infection of the liver and providing sterilizing immunity^2^. However, significant challenges exist in achieving sterilizing immunity against malaria. The rapid invasion of hepatocytes by sporozoites restricts the amount of time for antibodies to act; an effective vaccine must elicit sustained, high-titer antibody responses.The presence of even a single infected hepatocyte can initiate the blood stage, setting a high barrier for protection by pre-erythrocytic vaccine candidates^7^. Thus, innovative and multi-faceted vaccine strategies may be required to reproducibly provide strong protection.

*Anopheles* (*An*.) mosquitoes are the vectors of *Plasmodium* parasites, the causative agents of malaria. There are more than 400 species of *Anopheles*, approximately 70 of which are capable of transmitting human-infecting parasites, and 41 are considered dominant *Plasmodium* vector species^8,9^. *An. gambiae* has historically been the principal vector of *P. falciparum* in Africa, but beginning in 2013 Africa has seen an influx of *An. stephensi* mosquitoes, a species native to South Asia and parts of the Arabian peninsula^10^. *An. stephensi* are efficient vectors for both *P. falciparum* and *P. vivax*, and unlike other malaria vectors, are known to inhabit primarily urban environments. This raises the concern that malaria outbreaks within heavily populated urban areas in Africa will increase, even during the dry season, which could potentially lead to year-round malaria transmission^11^. *An. stephensi* have been firmly established in the Horn of Africa, and have spread further into surrounding countries including Kenya, Ghana, and Nigeria^3,12^.

Malaria begins with a bite from an infected mosquito. *Plasmodium* sporozoites, the infectious stage, are deposited into the dermis, epidermis, and bloodstream and then travel to the liver to seed the next stages of the parasite’s life cycle. Susceptibility to infection is influenced by a combination of host, pathogen, and environmental factors, one being the host response to mosquito saliva. Mosquitoes inject pathogens amidst their saliva when probing for a blood meal, and the vector-derived factors within the saliva have been shown to enhance infection, both in malaria and other vector-borne diseases. Multiple studies have shown that vector salivary components enhance infection across many vector species (*e*.*g*., mosquitoes^13–16^, ticks^17–19^, sand flies^20–23^, although their role in malaria remains unclear. Taken together, these studies suggest that inhibiting the activity of insect salivary proteins may be a useful strategy for decreasing the efficiency of vector-borne infections.

Although the role of mosquito saliva in the transmission of *Plasmodium* is somewhat controversial, recent studies have shown that the *Anopheles* salivary protein TRIO, expressed exclusively in female mosquitoes’ salivary glands and upregulated in infected mosquitoes^24,25^, influences the local inflammatory response in infected hosts, enhancing *Plasmodium* motility and infection^26^. Correspondingly, immunization with full-length *An. gambiae* TRIO (AgTRIO) protein or passive immunization with an AgTRIO-specific monoclonal antibody (mAb 13F-1) results in a reduction in *Plasmodium* liver burden in mice^25,26^. These data suggest that vaccines that target TRIO could hinder sporozoite migration and limit infection, especially if used in combination with a *Plasmodium*-specific target. Here we investigate the efficacy of bacteriophage virus-like particle (VLP)-based vaccines targeting epitopes from the *Anopheles* salivary protein, TRIO. Additionally, we investigate the efficacy of a combination vaccine consisting of TRIO VLPs and VLPs targeting the highly vulnerable L9 epitope from the *P. falciparum* circumsporozoite protein^27^. TRIO-epitope displaying VLPs elicited high-titer and long-lasting TRIO-reactive antibody responses that did not drop for ∼18 months post-immunization, essentially the lifespan of the mouse. In a mouse challenge model, immunization with TRIO VLPs resulted in a modest reduction in liver parasitemia that was more pronounced in mice that received a lower parasite challenge dose. However, co-immunization of TRIO VLPs did not enhance the protection provided by the CSP-targeted L9 VLP.

## METHODS

### Ethics statement for animal studies

All animal research complied with the Institutional Animal Care and Use Committee at the University of New Mexico School of Medicine (approved protocol #: 22-201289-HSC) and at Johns Hopkins University (approved permit #: MO18H419).

### Expression and purification of bacteriophage Qβ VLPs

Qβ bacteriophage VLPs were produced and purified as previously described^27^. Briefly, Qβ bacteriophage coat protein was expressed from the plasmid pETQCT in electrocompetent *E*.*coli* C41(DE3) cells (Sigma-Aldrich). Bacterial pellets were resuspended in lysis buffer (100mM NaCl, 10mM EDTA, 50mM Tris-HCl, 0.45% deoxycholate) and incubated on ice for 30min, followed by 3-5 cycles of sonication at 20Hz, until the solution was clear. Following sonication, residual DNA was removed using 10µg/mL DNase, 2.5mM MgCl_2_, and 0.05mM CaCl_2_ (all final concentrations) upon incubation on ice for 1 hour. The supernatant was isolated by centrifugation at 10,000rpm for 30min (TA-14-50 fixed-angle rotor). Ammonium sulfate was added to the supernatant to make a 70% solution and incubated on ice for 15min. Following incubation, precipitated protein was spun at 10,000rpm for 15min and the pellet was resuspended in cold SCB buffer (10mM Tris-HCl, 100mM NaCl, 0.1mM MgSO_4_). The solution was fractionated by size exclusion chromatography on a Sepharose CL-4B column. Fractions that contained Qβ VLPs were identified by gel electrophoresis and incubated in 70% ammonium sulfate overnight at 4°C to precipitate out protein. Following centrifugation at 10,000rpm for 15min, pellets were resuspended in PBS (pH 7.4) and dialyzed twice against PBS (pH 7.4). Prior to peptide modification, Qβ VLPs were depleted of endotoxin by 4 rounds of phase separation using Triton X-114 (Sigma-Aldrich)^28^. The final concentration of Qβ VLPs was determined via SDS-PAGE using known concentrations of hen’s egg lysozyme as a control

### Conjugation of peptides to Qβ VLPs

The L9 epitope of CSP was synthesized (GenScript) with a C-terminal linker sequence *gly-gly-gly-cys* (NANPNVDPNANPNVD-*GGGC*). Two TRIO peptides, derived from *An. gambiae* TRIO (VDDLMAKFN-*GGGC*) and *An. stephensi* TRIO (AANLRDKFN-*GGGC*), were synthesized (GenScript) with the same linker sequence. Each peptide was conjugated separately to the exposed surface lysine residues on Qβ VLPs using the heterobifunctional amine-to-sulfhydryl crosslinker, succinyl 6-[(β-maleimidopropionamido)hexanoate] (SMPH; ThermoFisher Scientific). SMPH was incubated with Qβ VLPs at a molar ratio of 10:1 (SMPH:Qβ coat protein) for 2 hours at room temperature. Excess SMPH was removed using an Amicon Ultra-4 centrifugal unit with a 100 kDa cutoff (Millipore). Peptides were individually added to Qβ VLPs at a molar ratio of 10:1 (peptide:Qβ coat protein) and incubated overnight at 4°C. Conjugation efficiency was measured by SDS-PAGE, where peptide addition can be seen by a shift in molecular weight. The percentage of coat protein with zero, one, two, or more attached peptides was determined by SDS-PAGE and used to calculate average peptide density per VLP.

### Expression and characterization of TRIO proteins

N-terminally His-tagged AgTRIO and AsTRIO, without the signal peptide, were synthesized and cloned into a pET-28a(+) vector (Twist Bioscience) and expressed using electrocompetent C41(DE3) *E. coli* cells (Sigma). Expression was induced by addition of 1mM IPTG at 37°C for 3 hours. Recombinant protein was purified using Ni-NTA resin (ThermoFisher Scientific). Western blot was used to evaluate protein content. After protein transfer, nitrocellulose membranes were blocked with PBS-T (PBS with 0.1% Tween 20) with 5% milk for 1 hour at room temperature. Membranes were blotted with sera from immunized mice or with an anti-6xHis-tag antibody (Cell Signaling Technology) as a positive control. Blots were stained with 1:4000 dilutions of HRP-labeled secondary antibodies and detected using Pierce ECL Western Blotting Substrate (ThermoFisher Scientific). Blots were imaged using a ChemiDoc MP Imaging System (BioRad).

### Mouse immunization studies

Groups of 4–6-week-old female Balb/c mice (Jackson Laboratory) were immunized intramuscularly with 5µg L9 VLPs, TRIO VLPs, or a combination of the two (5µg of each), for a total of 2 doses 3 weeks apart (n=5 per group). Control groups received 5µg unmodified wild-type (WT) VLPs. Serum was collected after each immunization and then periodically for up to 18 months following the initial immunization.

### Quantitating antibody responses

Serum antibodies against full-length CSP, TRIO peptides, and full-length TRIO were detected by ELISA. Recombinant CSP was expressed in *Pseudomonas fluorescens* and was generously provided by Gabriel Gutierrez (Leidos, Inc.)^29^. For the full-length protein ELISAs, wells of Immulon 2 ELISA plates (ThermoFisher Scientific) were coated with 250ng protein in 50µL 1xPBS and incubated overnight at 4°C. Wells were blocked with PBS-0.5% milk for 1 hour at room temperature. Sera isolated from immunized mice were serially diluted in PBS-0.5% milk and applied to wells overnight at 4°C. Reactivity was measured by addition of HRP-labeled goat anti-mouse IgG, diluted 1:4000 in PBS-0.5% milk (Jackson Immunoresearch), and detected by addition of TMB substrate. Reactions were stopped using 1% HCl and Optical Densities (ODs) were measured at 450nm.

Peptide ELISAs require an additional coating step. ELISA plates were initially coated with 500ng streptavidin at 4°C overnight. SMPH was added at 250ng/well and incubated for 1 hour at room temperature. Wells were coated with 250ng/well of peptide and incubated for 2 hours at room temperature. Wells were then blocked, incubated with serial dilutions of immunized mouse sera, and detected as described above.

### Mouse Pb-PfCSP-Luc sporozoite challenge

Challenge studies were performed using female 6–8-week-old C57BL/6 mice (n=5-7 per group). Mice were immunized intramuscularly with 5µg VLPs with or without adjuvant at 3-week intervals, for a total of 3 doses. The combination vaccine group received 5µg L9 VLPs and 5µg TRIO VLPs. All experimental groups received either 100µg Advax-3 (generously provided by Vaxine) or 2µg Cquim adjuvant (generously provided by ViroVax). Control groups included naïve mice and mice that were immunized with unmodified WT VLPs. Each immunogen was blinded to minimize the potential for bias in animal handling during the challenge portion of the study. Serum was collected following the third immunization.

Mice were challenged using transgenic *P. berghei* sporozoites engineered to express luciferase and full-length *P. falciparum* CSP in place of *P. berghei* CSP, denoted as Pb-PfCSP-luc. Mice were challenged by infected mosquitoes. *A. stephensi* mosquitoes were infected by blood feeding on Pb-PfCSP-luc sporozoite infected mice. Prior to challenge, mice were anesthetized with 2% Avertin and exposed to five infected mosquitoes. Mosquitoes that had taken a blood meal were counted. Liver burden was measured 42 hours post-challenge by intraperitoneally injecting mice with 100µL ⫐-luciferin (30mg/mL) and measuring liver luminescence using an IVIS Spectrum Imaging System (Perkin Elmer). Five days later, blood smears were evaluated by Giemsa staining for parasitemia.

### Statistical analysis

All statistical analyses were performed using GraphPad Prism v.10. Two-sided t-tests were used for immunogenicity studies and Mann-Whitney tests were used for the mosquito challenge experiments. For percent inhibition calculations, liver luminescence values of individual vaccinated mice were divided by the mean of mice in negative control groups. Background luminescence levels were subtracted from all values.

## RESULTS

### Construction and characterization of TRIO VLPs

To generate a TRIO-specific vaccine, we targeted the 13F-1 epitope, a 9 amino acid peptide near the C-terminus of AgTRIO (VDDLMAKFN; Fig. 1A) that was previously identified by Chuang and colleagues^26^. The 13F-1 mAb was shown to recognize TRIO in salivary gland extracts from both *An. gambiae* and *An. stephensi*, although it is only weakly reactive with *An. stephensi* extracts^26^ which is unsurprising because only 4/9 amino acids in the 13F-1 epitope are shared between AgTRIO and *An. stephensi* TRIO (AsTRIO) (Fig.1A). Therefore, we opted to engineer two separate bacteriophage VLPs displaying the AgTRIO peptide and the homologous AsTRIO peptide (AANLRDKFN; Fig. 1A) at high valency. AgTRIO and AsTRIO peptides were synthesized with a C-terminal linker sequence (-Gly-Gly-Gly-Cys). Peptides were then conjugated to the surface of Qβ bacteriophage VLPs, as previously described^30^, through a bifunctional cross-linker (SMPH) to produce AgTRIO and AsTRIO VLPs (Fig. 1B). Peptide-VLP conjugation efficiency was measured by SDS-PAGE. Fig. 1C, lane 3, shows that virtually all of the Qβ bacteriophage coat protein subunits display the AgTRIO peptide. Fig. 1C, lane 4, shows that more than half of the Qβ bacteriophage coat protein subunits display the AsTRIO peptide, based on the disappearance of the 14kDa band that represents unbound Qβ coat protein. We estimate that >300 copies of AgTRIO and >200 copies of AsTRIO were conjugated to each VLP.

**Figure 1:**
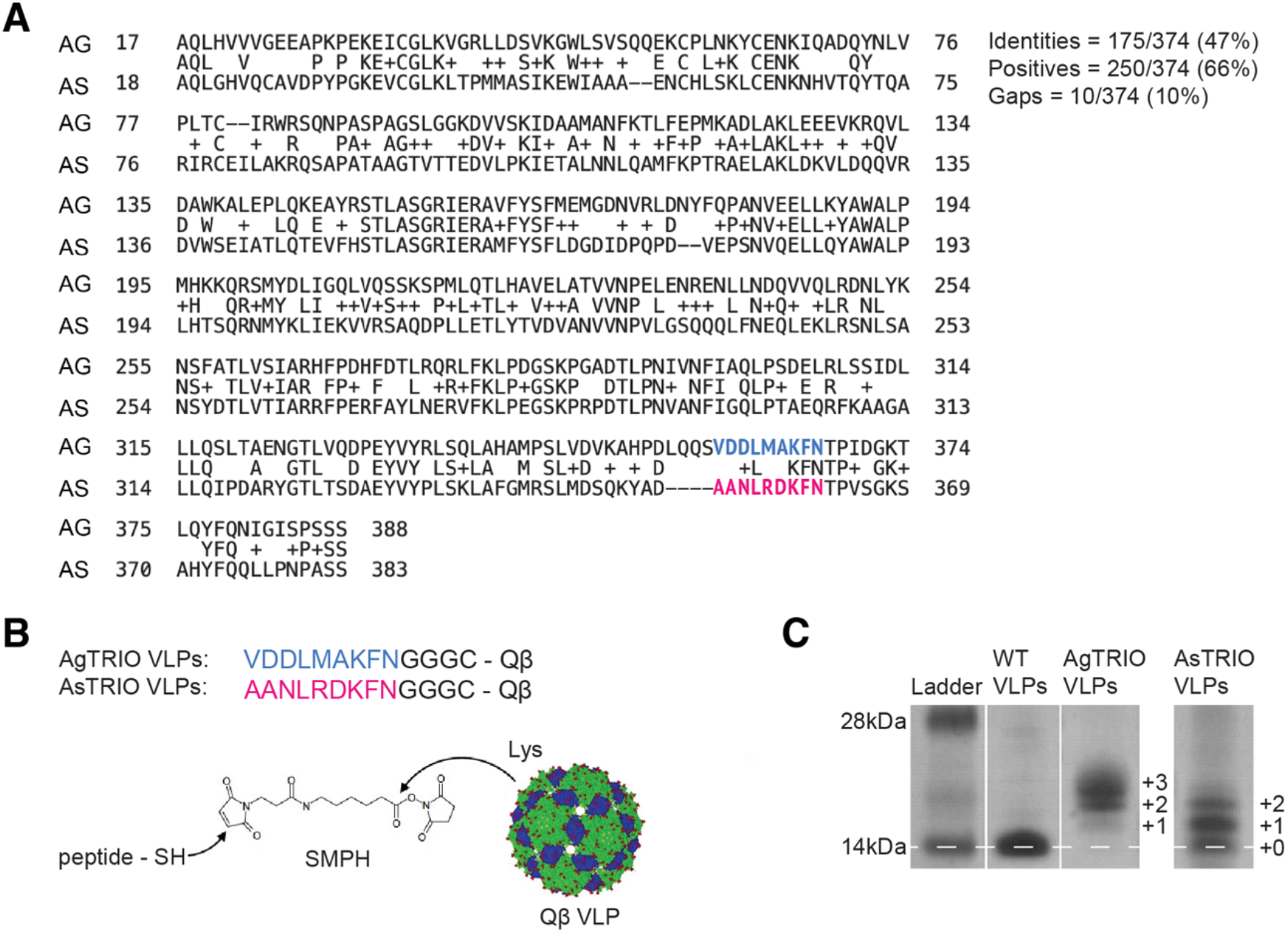
TRIO VLP design and characterization. (A) Amino acid sequences of the salivary protein TRIO from *Anopheles gambiae* (AgTRIO) and *Anopheles stephensi* (AsTRIO). Peptides used in vaccines are highlighted blue (AgTRIO) and magenta (AsTRIO). Multiple sequence alignment was produced using NCBI BLASTp suite (AgTRIO accession number AAL68795.1). (B) AgTRIO and AsTRIO peptides are synthesized with a C-terminal linker sequence (-Gly-Gly-Gly-Cys) which binds to the maleimide arm of the chemical cross-linker, SMPH. The NHS-ester arm of SMPH subsequently binds to surface exposed lysines on the Qβ bacteriophage VLP. (C) SDS-PAGE analysis of peptide-conjugated VLPs. Unmodified Qβ bacteriophage coat protein found in WT VLPs (lane 2, no peptide attached) has a molecular weight of 14kDa. Conjugation efficiency is assessed via shifts in molecular weight, based on the number of peptides attached per coat protein. Gel images are from the same gel.

### AgTRIO and AsTRIO VLPs induce high-titer antibody responses which bind to full-length TRIO

To assess the immunogenicity of the AgTRIO and AsTRIO VLPs, mice were immunized intramuscularly with 5µg AgTRIO VLPs or 5µg AsTRIO VLPs and boosted with the same dose three weeks later. One week following the boost, sera were collected and antibody responses against the targeted TRIO peptide epitopes or full-length TRIO protein were measured by ELISA. To measure responses against full-length TRIO proteins, we expressed and purified recombinant full-length AgTRIO and AsTRIO proteins in *E. coli* (Fig. 2A,B). Serum antibody levels against the AgTRIO and AsTRIO peptides (Fig. 2C,D) and full-length TRIO proteins (Fig. 2E,F) were measured by end-point dilution ELISA. We measured antibody responses against the targeted protein as well as cross-species reactivity. Immunization with AgTRIO VLPs resulted in high IgG antibody titers against both the AgTRIO peptide (Fig. 2C) and full-length AgTRIO (Fig. 2E). Immunization with AsTRIO VLPs similarly resulted in high titers against the AsTRIO peptide (Fig. 2D) and full-length AsTRIO protein (Fig. 2F), although titers against full-length AsTRIO were slightly lower relative to AgTRIO responses. This result may reflect the challenges we encountered in producing full-length AsTRIO; preparations of AsTRIO were less pure and less concentrated than AgTRIO. Importantly, negative control sera from mice immunized with VLPs bound to a different unrelated *Aedes aegypti* salivary peptide (non-specific VLPs) did not show reactivity to full-length AgTRIO or AsTRIO or to the peptides. We also observed some cross-species recognition of full-length TRIO; AsTRIO VLPs recognize full-length AgTRIO (Fig. 2E, magenta), and AgTRIO VLPs recognize full-length AsTRIO (Fig. 2F, blue), likely mediated by antibodies that are directed to the conserved C-terminal domain of the TRIO epitope. These results indicate that immunization with either AgTRIO or AsTRIO VLPs could offer some cross-species reactivity in an area with multiple *Anopheles* vector species.

**Figure 2:**
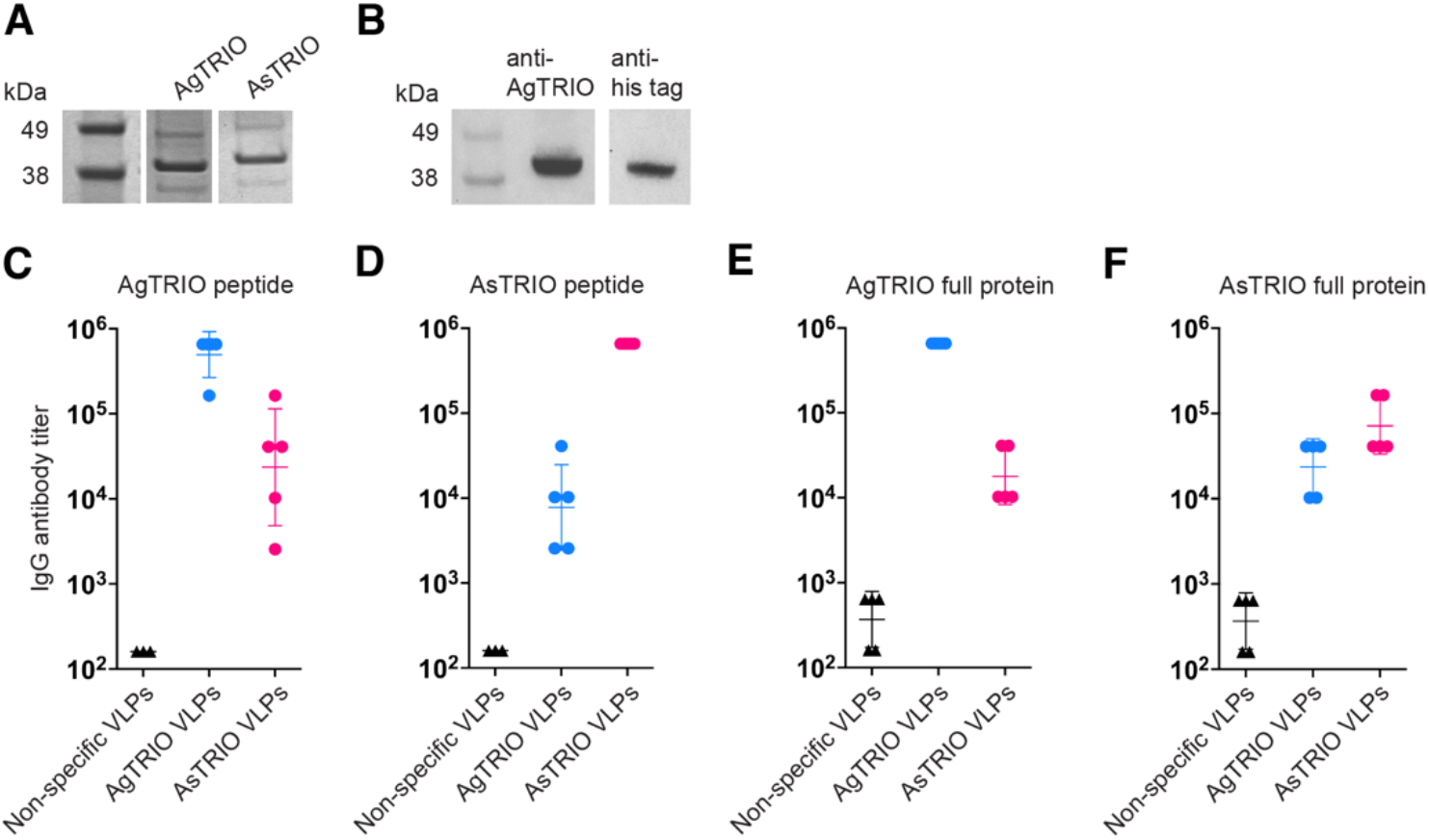
Immunization with TRIO peptide VLPs results in high-titer antibody responses against peptides and full-length TRIO proteins. Full-length AgTRIO and AsTRIO proteins containing N-terminal 6x-His tags were expressed in *E. coli* and purified by Ni-NTA resin. (A) SDS PAGE of full-length AgTRIO and AsTRIO proteins before Ni-NTA purification. Both proteins are approximately 40kDa. (B) Western blot membranes containing 5µg of the purified AgTRIO protein were blotted with sera from AgTRIO VLP immunized mice (anti-AgTRIO, lane 2) or anti-6x his tag antibody (anti-his tag, lane 3). Blots are from the same membrane, imaged using a ChemiDoc MP Imaging system (BioRad). (C-F) Mice (n=5) were immunized with 5ug of either AgTRIO (blue) or AsTRIO peptide (magenta) VLPs and boosted 3 weeks later. End-point dilution IgG antibody titers were measured 1 week after the boost by ELISA against (C) AgTRIO peptide, (D) AsTRIO peptide, (E) full-length AgTRIO, and (F) full-length AsTRIO. Sera from mice immunized with an unrelated mosquito salivary peptide were used as a negative control (non-specific VLPs; 1-2 weeks after the boost). Data is reported as geometric mean ± geometric SD.

### Combination of TRIO VLPs with VLPs targeting the *P. falciparum* circumsporozoite protein elicits durable, high-titer antibody responses

The majority of pre-erythrocytic vaccines against *P. falciparum* target the circumsporozoite protein (CSP), a protein which is abundantly expressed on the surface of the parasite and is involved in motility and hepatocyte invasion^31^. The recently approved malaria vaccines, RTS,S/AS01 and R21/Matrix-M, target the C-terminal half of CSP, a region that contains the major repeat domain and the C-terminal domain^5,6^. However, there is emerging evidence that antibodies that target epitopes in the junctional region of CSP, which is located at the N-terminus of the central repeat region that is not included in either vaccine, can confer stronger protection from infection. For example, passive immunization the monoclonal antibody L9, which targets an epitope consisting of alternating minor (NVDP) and major (NANP) repeat sequences, strongly protects both mice and humans from malaria infection^32,33^. We developed a VLP-based vaccine that targets the L9 epitope and showed that it could provide sterilizing immunity in a mouse malaria challenge model^27^. Previous studies have shown that passive immunization with a combination of anti-CSP and anti-TRIO antibodies results in decreased *P. berghei* liver burden in mice^25^. Therefore, we were interested in assessing the levels and durability of antibody responses as well as protection mediated by a combination vaccine consisting of VLPs displaying TRIO with VLPs displaying the L9 epitope from CSP.

Mice were immunized with a combination vaccine consisting of 5µg of L9 VLPs and 5µg of TRIO VLPs (10µg VLPs total; Table 1). Mice were immunized intramuscularly with L9 and AgTRIO VLPs (Fig. 3A) or L9 and AsTRIO VLPs (Fig. 3B) and boosted 3 weeks later. Anti-CSP and anti-TRIO peptide titers were measured to assess the immunogenicity of the L9 and TRIO VLPs, respectively. Titers were measured out to ∼18 months for the L9 and AgTRIO VLP combination, and out to 1 year for the L9 and AsTRIO VLP combination. Importantly, both components of the combination vaccines elicited similar IgG antibody titers; thus, it appears that the response to one VLP does not lower the antibody titers against the other. Moreover, antibody responses against both target antigens were exceptionally durable. Although there was a slight decline in antibody levels over the course of the study, these results demonstrate, and confirm our results from previous work^27,34–36^ demonstrating that VLPs consistently elicit high-titer, long-lasting antibodies against multivalently displayed peptides.

**Table 1:**
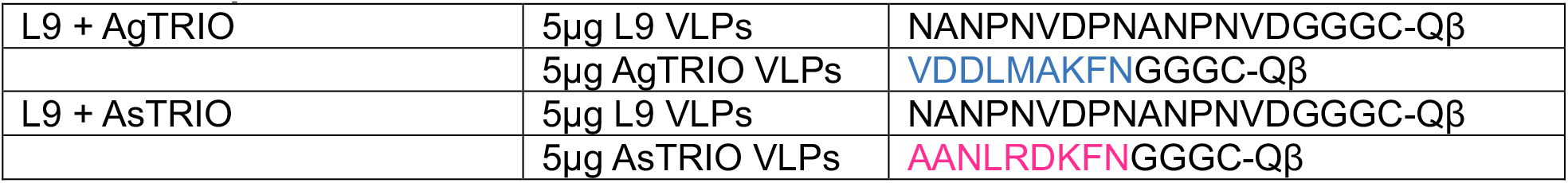
Composition of combination vaccines.

|  |  |  |
| --- | --- | --- |
| L9 + AgTRIO | 5µg L9 VLPs | NANPNVDPNANPNVDGGGC-Qβ |
|  | 5µg AgTRIO VLPs | VDDLMAKFNGGGC-Qβ |
| L9 + AsTRIO | 5µg L9 VLPs | NANPNVDPNANPNVDGGGC-Qβ |
|  | 5µg AsTRIO VLPs | AANLRDKFNGGGC-Qβ |

**Figure 3:**
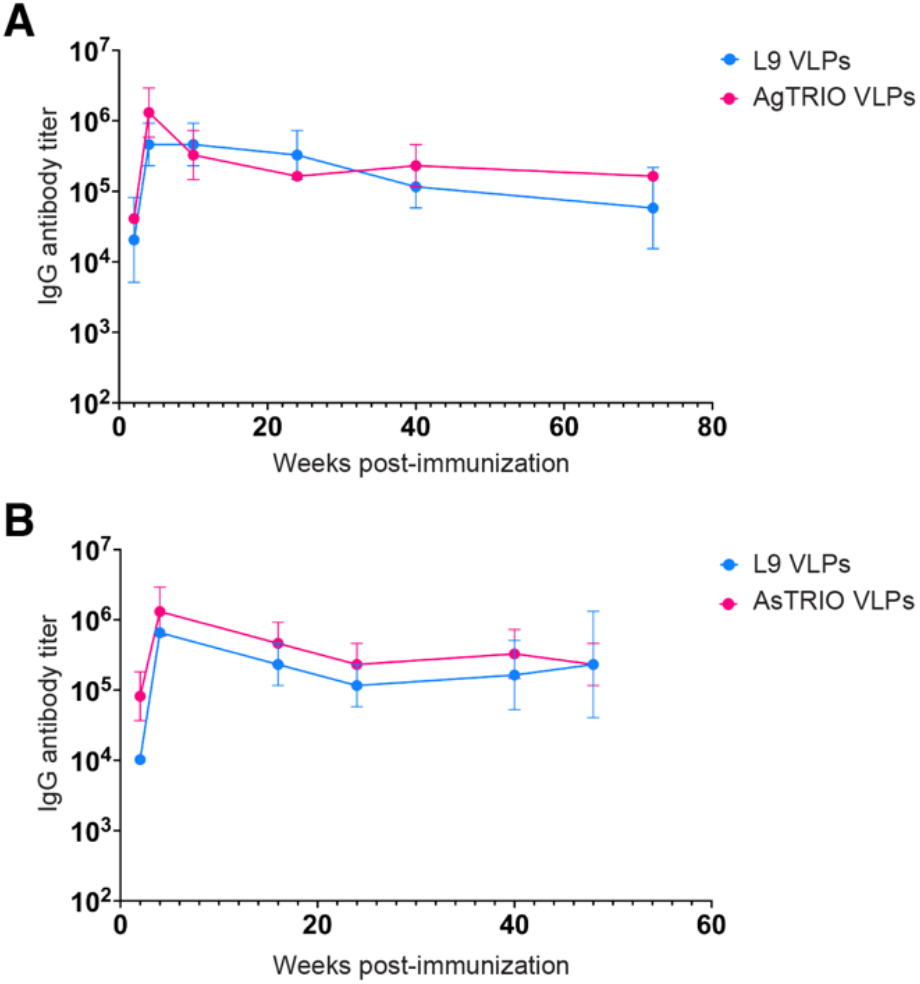
Immunization with a combination of L9 and TRIO VLPs results in durable, high-titer antibody responses. Mice (n=5) were immunized with 5µg of L9 VLPs and 5µg of TRIO VLPs and boosted 3 weeks later. IgG antibody titers were measured by ELISA against TRIO peptides (magenta) and full-length CSP (blue) from *Plasmodium falciparum*. (A) IgG titers from mice immunized with a combination of L9 VLPs and AgTRIO VLPs were measured out to 18 months post-immunization. (B) IgG titers from mice immunized with a combination of L9 VLPs and AsTRIO VLPs were measured out to 1-year post-immunization. Data is reported as geometric mean ± geometric SD.

### Immunization with TRIO VLPs alone does not strongly protect mice from *Plasmodium*challenge

To evaluate protection mediated by TRIO VLPs alone or the combination vaccine (L9 and TRIO VLPs), mice were immunized three times and tested in a well-established mouse model for evaluating CSP-targeted pre-erythrocytic vaccines^37^. Mice were challenged with mosquitos carrying transgenic *P. berghei* sporozoites that contain CSP from *P. falciparum* and also express a luciferase reporter (*Pb-PfCSP-Luc* sporozoites) that allows the quantification of parasite liver burden. Because *An. stephensi* mosquitoes were used in the challenge experiments, mice were vaccinated with AsTRIO VLPs. Female C57BL/6 mice were challenged with five infected mosquitoes to more closely recapitulate natural infection. A caveat to this method is that the amount of sporozoites in each mosquito and the amount delivered via bite cannot be controlled, resulting in some variability in the parasite liver burden between experiments.

We have previously shown that adjuvants can increase anti-CSP antibody responses elicited by VLP-based vaccines and that high-titer antibodies are critical for the efficacy of CSP-targeted VLP-based vaccines^27^. Thus, in the challenge experiments, we evaluated the ability of two different adjuvants to increase the immunogenicity of L9/TRIO vaccines in two separate experiments. In the first challenge experiment vaccines were combined with Advax-3 adjuvant (which is a mixture of CpG55.2 oligonucleotide, a TLR9 agonist, with aluminum hydroxide), and in the second challenge experiment vaccines were combined with Carbomeriquim (Cquim) adjuvant (a dual TLR7/8 agonist). Prior to challenge, anti-AsTRIO and anti-CSP IgG titers in C57Bl/6 mice immunized with AsTRIO VLPs alone or with the combination vaccine were evaluated by ELISA (Fig. 4D,G). There were no significant differences in anti-AsTRIO IgG titers between the groups and between the two challenges. Comparing the groups that received the combination vaccine, anti-CSP antibody levels were higher in the group that received Cquim adjuvant.

**Figure 4:**
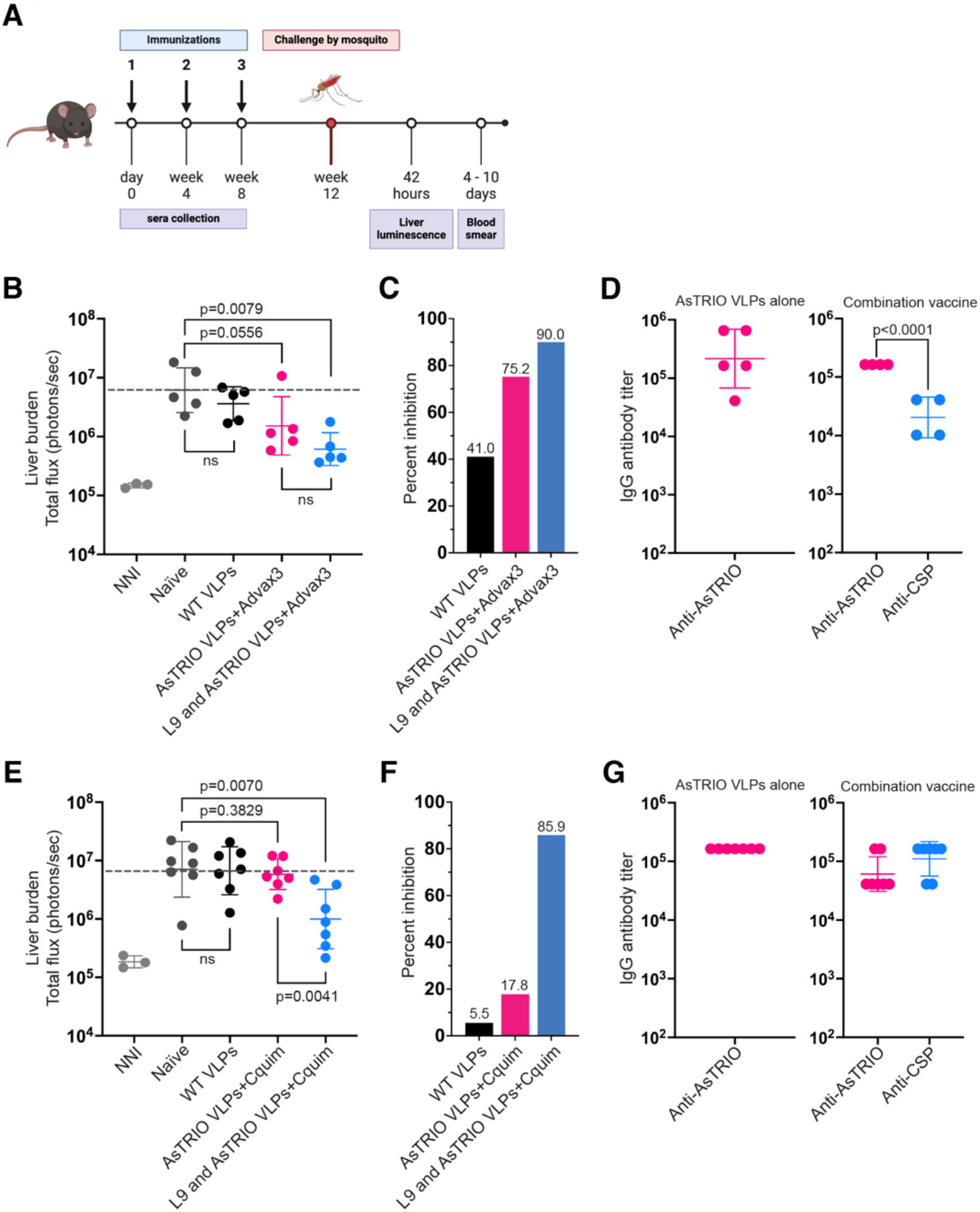
Immunization with TRIO VLPs alone is not sufficient to prevent liver-stage infection. Female C57BL/6 mice were immunized with 5µg doses of WT VLPs (negative control), AsTRIO VLPs, or a combination of L9 and AsTRIO VLPs (5µg of each) and boosted 3 and 6 weeks later. Two different adjuvants were included in specified groups (Advax-3 and Cquim). (A) Outline of immunization and challenge schedule. Groups of C57BL/6 mice (n = 5-7 per group) were immunized three times and then challenged with five *Pb-PfCSP-Luc* infected mosquitos. (B) and (C) show data from the first challenge experiment, and (E) and (F) are from a second challenge experiment. (B,E) Parasite liver burden was measured by luminescence 42 hours after mosquito challenge. The grey dotted line marks the mean liver burden of mice immunized with WT VLPs. Background luminescence was determined using three uninfected mice (NNI, naïve non-infected). (C, F) Percent inhibition of liver infection calculated from luminescence data (relative to the mean signal in the negative control groups). (D, G) Endpoint dilution IgG titers were measured by ELISA on sera collected 6 weeks after primary immunization (1 week prior to mosquito challenge). Data is reported as geometric mean ± geometric SD. P value was determined using a two-tailed Mann-Whitney test for (B, C, E, F) and a two-tailed t test for (D, G). (A) was created with BioRender.com.

In the first challenge (using Advax-3 adjuvant), both immunization with AsTRIO VLPs alone and the combination of AsTRIO and L9 VLPs resulted in reduced liver parasite burden compared to naïve controls (Fig. 4B). Immunization with AsTRIO VLPs alone resulted in a 75.2% reduction (p=0.0556) in liver parasite burden compared to naïve challenged mice (Fig. 4C). Immunization with the combination TRIO/L9 vaccine reduced parasite liver loads by 90% (p=0.0079) (Fig. 4B, C), which is similar to what we previously reported using the Advax-3 adjuvanted L9 VLPs alone^27^. These data indicate that immunization with AgTRIO VLPs can reduce parasite liver burden, but that AgTRIO VLPs do not synergize with L9 VLPs to enhance protection. However, protection from liver infection in the group immunized with AsTRIO VLPs was less apparent in the second challenge experiment. In this study, immunization with AsTRIO only resulted in a modest 17% reduction in parasite liver burden, and this difference was not statistically significant (p=0.38) (Fig. 4E,F). In both experiments, all mice immunized with AsTRIO VLPs developed blood parasitemia by day 5 post-challenge. One caveat to the second challenge experiment is that the mean parasite liver burden was higher across all groups. Thus, it is possible that antibodies against TRIO may have some protective efficacy at lower challenge doses. Overall, these data indicate immunization with AsTRIO VLPs may provide some benefit in decreasing parasite load in the liver, but these effects are subtle and do not lead to sterilizing immunity. Further research into the mechanism of protection is required, especially on how AsTRIO VLPs affect sporozoite motility and dispersal in the dermis; it has been shown that AgTRIO antiserum significantly decreases sporozoite dispersal in the dermis^25^, but not much is known about the mechanism of action.

## DISCUSSION

The initial stage of malaria infection in which sporozoites, the infectious form of the malaria parasite, must transit through the epidermis and dermis prior to establishing liver infection is a potential period of vulnerability that could be targeted using vaccines. It was shown that antibodies targeting *Plasmodium* sporozoites can exert their protective effect at the bite site by impacting the mobility and migration of sporozoites^38^. To complement the role of anti-sporozoite antibodies, we designed a virus-like particle (VLP) vaccine targeting TRIO, a salivary protein found in many *Anopheles* species that are vectors for human malaria parasites, including *An. gambiae, An. arabiensis, An. stephensi, and An. albimanus*. It is one of several salivary proteins that are overexpressed in salivary glands infected with *Plasmodium*^39,40^. Previous work from Fikrig and colleagues has demonstrated that antibodies against the *An. gambiae* TRIO protein diminish sporozoite speed and migration in murine dermis^25^. Further, they identified a monoclonal antibody against a linear epitope within the C-terminal region of the TRIO protein that was able to reduce early *Plasmodium* infection in mice^26^. For these reasons, we decided to test the efficacy of a VLP-based vaccine targeting this TRIO epitope in reducing *Plasmodium* liver burden. Additionally, we investigated if it acts synergistically with a VLP-based vaccine that targets the *P. falciparum* circumsporozoite protein, which we previously showed could provide sterilizing immunity in approximately 60% of mice^27^.

TRIO VLPs induced high-titer antibodies that were extremely durable; titers did not drop significantly for up to 18 months, essentially the lifespan of a mouse (Fig. 3). We showed that *An. gambiae* TRIO VLPs (AgTRIO) can recognize TRIO derived from *An. stephensi* (AsTRIO), and vice versa (Fig. 2). But, based on the lower cross-species antibody titers and sequence heterogeneity of TRIO proteins, we elected to target each TRIO protein individually.

Although TRIO VLPs were shown to be immunogenic and produce long-lasting antibody responses (Fig. 2 and 3), immunization with these VLPs only resulted in modest reductions in liver parasitemia in mice (Fig. 4). Moreover, our data indicate that VLPs displaying this specific epitope from TRIO do not significantly add to the protection provided by VLPs targeting the vulnerable L9 epitope from CSP. However, our data indicate that the efficacy of immunization with TRIO VLPs may be dependent on initial sporozoite burden, perhaps because the TRIO vaccine induces antibodies that do not directly target sporozoites themselves. Mice in these studies were infected by mosquito challenge, which makes it difficult to control the amount of sporozoites introduced. This is illustrated in Figure 4, (Fig. 4B and Fig. 4E), where challenge 2 has a higher overall mean parasite liver burden (∼2 times higher) than challenge 1, and this was mirrored in decreased protection provided by both the TRIO VLP vaccine and the combination vaccine. On one hand, this does mimic the variation one would see in a natural infection^41^, on the other, humans would typically receive much less than in an experimental setting; humans typically receive fewer than 50 sporozoites per bite on average^42^. The TRIO vaccine may be more effective in a situation where the number of mosquitoes in the challenge are reduced or the number of sporozoites is controlled. It may be valuable to re-evaluate the ability of the TRIO vaccine to protect against different challenge doses.

## ACKNOWLEDGEMENTS

We thank Sunil David and ViroVax, LLC for generously providing Cquim adjuvant.

